# The effect of resource depletion on the thermal response of mosquito population fitness

**DOI:** 10.1101/2021.02.12.430918

**Authors:** Paul J. Huxley, Kris A. Murray, Lauren J. Cator, Samraat Pawar

## Abstract

The temperature-dependencies of life history traits are increasingly being used to predict how climatic warming will affect vector-borne disease dynamics, partially by affecting the abundance dynamics of the vector population. Such predictions generally arise from mathematical models that incorporate the temperature dependence of traits measured under laboratory conditions. These temperature-trait relationships are typically estimated from juvenile populations reared under optimal resource conditions, even though natural populations experience intermittent resource depletion. Using laboratory experiments on the mosquito *Aedes aegypti*, combined with a stage-structured population model, we show that resource depletion in the juvenile habitat can significantly depress the vector’s maximal population growth rate (*r*_m_) across the entire temperature range, cause it to peak at a lower temperature, and narrow its thermal niche width. Our results provide compelling evidence for future studies to consider resource depletion when predicting the effects of global change on vector-borne disease transmission, disease vectors and other arthropods.

## INTRODUCTION

Global environmental change is predicted to affect the spatiotemporal distributions of arthropod disease vectors and the diseases they transmit (Mordecai et al. 2020, WHO 2020). For example, recent studies suggest that climatic warming may increase the thermal suitability for Zika virus transmission, leading to 1.3 billion more people being at risk of exposure by 2050 (Ryan *et al.* 2021). Other studies have predicted that warming will increase the global invasion potential of *Aedes aegypti*, a principal vector of dengue, yellow fever and chikungunya (Iwamura *et al.* 2020). Such predictions often arise from disease transmission models that incorporate thermal performance curves (TPCs) for vector life history traits, such as juvenile development and mortality, which together define the TPC of maximal population growth rate (*r*_m_, a measure of population fitness; Savage et al. 2004).

Typically, such trait-level TPCs are obtained from data that come from juvenile populations reared under optimal food conditions in the laboratory (e.g. Shocket *et al.* 2020). However, in nature, organisms experience significant spatiotemporal variation in resource availability, including intermittent resource depletion (Ostfeld & Keesing 2000; Yang *et al.* 2008; Beltran *et al.* 2021). While recent studies have shown that resource availability can affect the *r*_m_ TPC in aquatic ectotherms (Orcutt & Porter 1984; Koussoroplis & Wacker 2016; Thomas *et al.* 2017), very studies have examined how the *r*_m_ TPC in terrestrial arthropods may be shaped by variation in resource availability (but see Huxley *et al.* 2020). Therefore, the question of whether and how temperature and resource depletion interact to modulate disease transmission together through effects on underlying vector traits remains open.

Temperature and resource availability are fundamentally linked at the physiological level, so studies on their combined effects could provide important mechanstic insights into how species will respond to climate change (Taheri *et al.* 2021). Temperature alters food assimilation and energy expenditure rates thus, governs energy requirements (Lemoine & Burkepile 2012). As per capita energy requirements intensify with increasing temperature, resource depletion rates and the strength of intra-specific competition should also increase. These combined effects are bound to compromise the growth, development, and survival of individuals. These trait-level effects are then expected to propagate through the stage-structured population dynamics to affect the shape of the *r*_m_ TPC (Amarasekare & Savage 2012; Huey & Kingsolver 2019). This is because *r*_m_ is essentially proportional to the difference between biomass gained through consumption and that lost to respiration and mortality (Savage *et al.* 2004). Resource limitation and competition would be expected to decrease *r*_m_ across temperatures, as they would both undermine biomass intake and elevate biomass loss.

Furthermore, if the rate of biomass loss increases faster than any increase in biomass gain with temperature, the thermal optimum of *r*_m_ (*T*_opt_) may also shift downwards (García-Carreras *et al.* 2018). For the same reason, the range of temperatures over which *r*_m_ is positive (the thermal niche width) may become narrower. As a result, the combined effects of climatic warming and decreased resource availability could contribute to the contraction of species range boundaries. This effect could simultaneously decrease the burden of vector-borne diseases and agricultural pests but increase the extinction risk of vulnerable species (Amarasekare 2019; Lehmann *et al.* 2020). Conversely, concurrent increases in temperature and resource availability with climatic warming could have the opposite effect by optimising *r*_m_ thus, promoting the invasion of tropical taxa into temperate habitats (Amarasekare & Simon 2020). This effect could further increase the huge socioeconomic cost of invasions by disease vectors, such as *Aedes* mosquitoes (Diagne *et al.* 2021).

Despite the potential for variation in resource availability to mediate the *r*_m_ TPC through its effects on underlying traits, most studies of the effects of temperature × resource interactions on terrestrial arthropods lack important features. For example, studies on mosquito vectors do not provide a mechanistic understanding of how temperature and resource availability can interact to influence underlying *r*_m_ traits (Couret *et al.* 2014; Barreaux *et al.* 2018), or the population-dynamic mechanisms are not understood by which temperature- and resource-driven changes in component *r*_m_ traits can affect the shape of the *r*_m_ TPC (Reisen *et al.* 1984; Tun-Lin *et al.* 2000; Farjana *et al.* 2012; Padmanabha *et al.* 2012), or both (Keirans & Fay 1968). Although such studies are informative, a more generalisable mechanistic understanding is needed to understand and predict the population-level effects of environmental change on disease vectors, and other arthropods.

Addressing this gap is particularly important for organisms with complex life histories, such as mosquitoes. For example, *Aedes* mosquitoes are expected to be mainly regulated by larval competition for limited resources (Southwood *et al.* 1972; Dye 1984), and have juvenile stages that are restricted to small, ephemeral aquatic habitats that are susceptible to infrequent resource inputs (Arrivillaga and Barrera 2004, Barrera et al. 2006, Yee and Juliano 2012). Here, we investigate the effects of resource depletion on the *r*_m_ TPC by exposing *Ae. aegypti* larvae to a realistic range of temperatures and resource concentration levels. We show that resource depletion can significantly change the shape of the *r*_m_ TPC through its effects on underlying traits in a predictable and general way.

## RESULTS

All trait responses varied significantly with temperature and resource level, with a significant interaction between the two environmental variables (figure 1, tables 1, 2).

**Figure 1.**
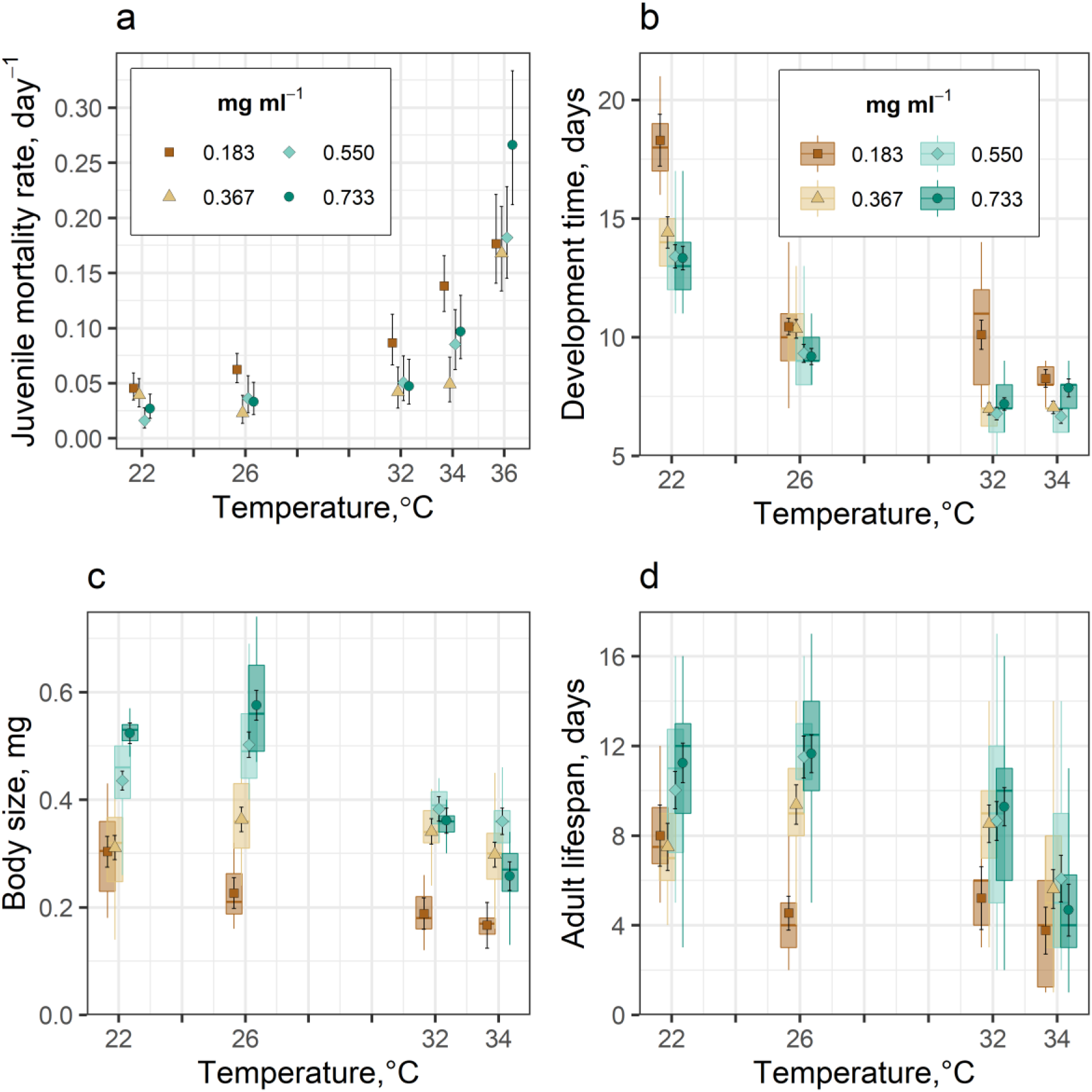
The effect of resource depletion on fitness traits in *Ae. aegypti* with 95% credible intervals (CIs). **a,** Resource depletion at low resource concentration (0.183 mg ml^−1^) increased the negative effect of increased temperature on juvenile mortality. **b**, Development time decreased with temperature at all resource levels but, at most temperatures, it was significantly extended by resource depletion at 0.183 mg ml^−1^. **c**, As temperatures increased from 22°C, resource depletion at 0.183 mg ml^−1^ significantly reduced size at emergence. **d**, As temperatures increased from 22 to 32°C, resource depletion at 0.183 mg ml^−1^ significantly reduced adult lifespan. Symbols denote the regression estimated means with 95% CIs calculated from the standard errors (replicate dropped, table 2) for the resource levels at each temperature. The resulting ANOVAs of the regressions for each trait are presented in table 1. Boxplot horizontal lines represent medians. Lower and upper hinges are the 25th and 75th percentiles. Upper whiskers extend from the hinge to the largest value no further than 1.5 × inter-quartile range (IQR) from the hinge. The lower whisker extends from the hinge to the smallest value at most 1.5 × IQR of the hinge.

**Table 1.**
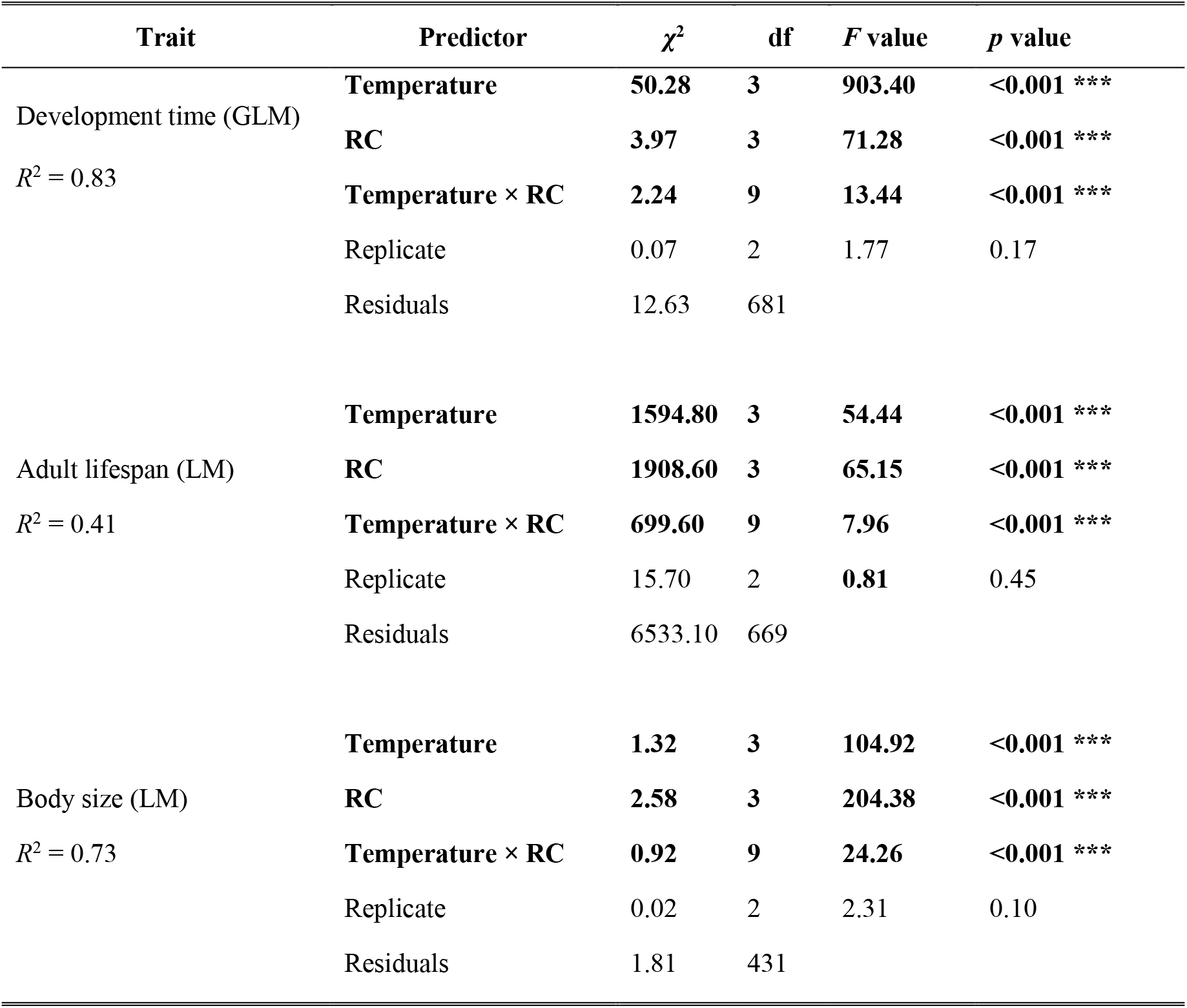
Type II Analysis of Variance results from regression models fitted to the responses of life history traits to temperature and resource concentration (RC) levels. Significant effects are shown in boldface type. * ⇒ *p* value<0.05; ** ⇒ *p* value<0.01 *** ⇒ *p* value<0.001.

| Trait | Predictor | $\chi^2$ | df | F value | p value |
| --- | --- | --- | --- | --- | --- |
| Development time (GLM)<br>$R^2 = 0.83$ | **Temperature** | **50.28** | **3** | **903.40** | **<0.001 \*\*\*** |
|  | **RC** | **3.97** | **3** | **71.28** | **<0.001 \*\*\*** |
|  | **Temperature × RC** | **2.24** | **9** | **13.44** | **<0.001 \*\*\*** |
|  | Replicate | 0.07 | 2 | 1.77 | 0.17 |
|  | Residuals | 12.63 | 681 |  |  |
| Adult lifespan (LM)<br>$R^2 = 0.41$ | **Temperature** | **1594.80** | **3** | **54.44** | **<0.001 \*\*\*** |
|  | **RC** | **1908.60** | **3** | **65.15** | **<0.001 \*\*\*** |
|  | **Temperature × RC** | **699.60** | **9** | **7.96** | **<0.001 \*\*\*** |
|  | Replicate | 15.70 | 2 | **0.81** | 0.45 |
|  | Residuals | 6533.10 | 669 |  |  |
| Body size (LM)<br>$R^2 = 0.73$ | **Temperature** | **1.32** | **3** | **104.92** | **<0.001 \*\*\*** |
|  | **RC** | **2.58** | **3** | **204.38** | **<0.001 \*\*\*** |
|  | **Temperature × RC** | **0.92** | **9** | **24.26** | **<0.001 \*\*\*** |
|  | Replicate | 0.02 | 2 | 2.31 | 0.10 |
|  | Residuals | 1.81 | 431 |  |  |

**Table 2.**
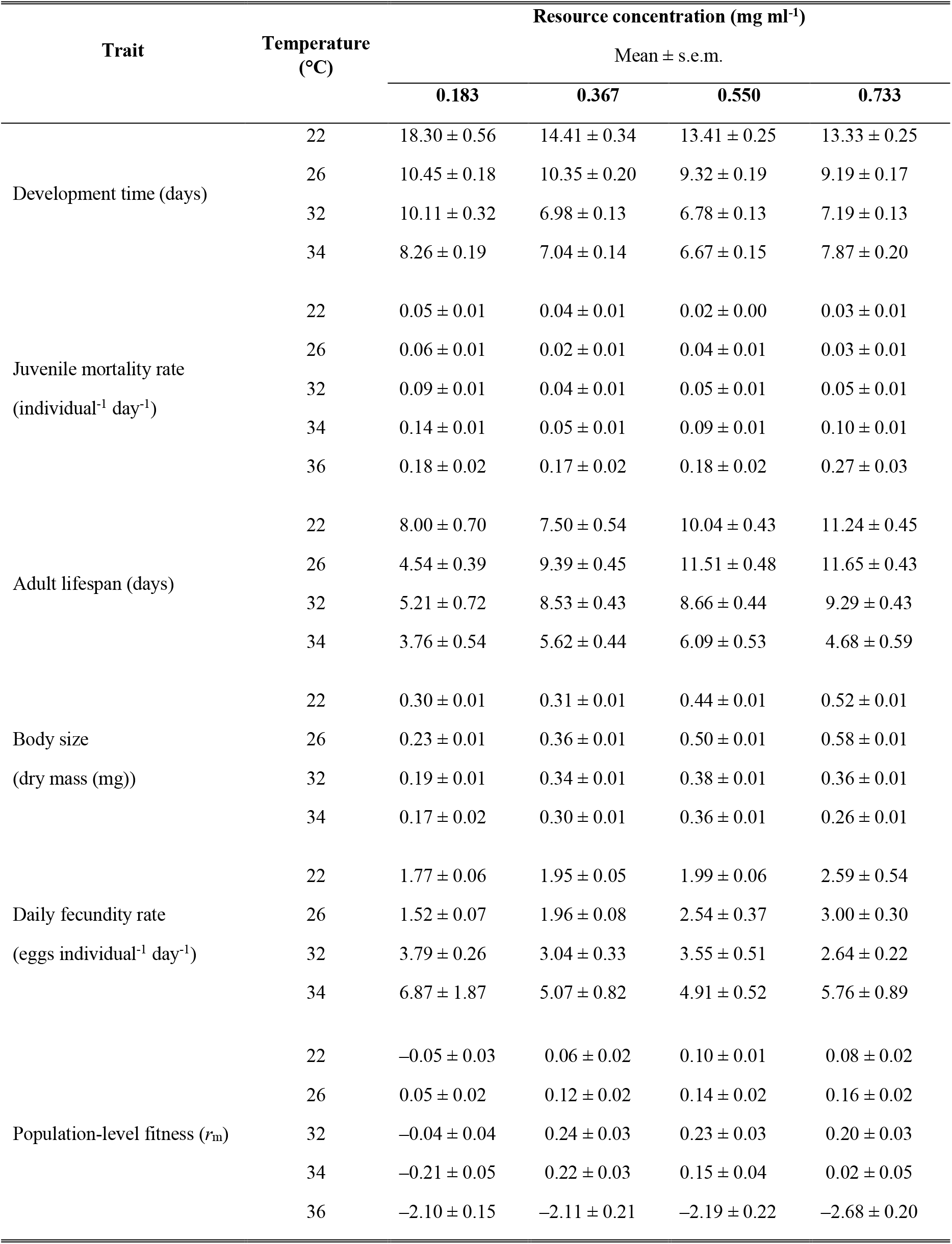
Comparison of the effect of resource depletion on the temperature-dependence of *r*_m_ and its component traits. The means with standard errors for juvenile mortality rate were estimated by fitting an exponential function to each treatment. The means with standard errors for development time, lifespan and size were estimated by using the statistical models in table 1 (replicate dropped). For fecundity, the standard errors were estimated using the `Rmisc` package in R. For *r*_m_, 95% CIs were approximated using the method described in Skalski *et al.* (2007). For *r*_m_ TPC fitting, non-positive matrix projection *r*_m_ values at 36°C were adjusted to ‒0.30. For plotting (figure 2a), non-positive *r*_m_ values were cut off at ‒0.10.

| Trait | Temperature<br>(°C) | Resource concentration (mg ml<sup>-1</sup>) |  |  |  |
| --- | --- | --- | --- | --- | --- |
|  |  | Mean ± s.e.m. |  |  |  |
|  |  | 0.183 | 0.367 | 0.550 | 0.733 |
| Development time (days) | 22 | 18.30 ± 0.56 | 14.41 ± 0.34 | 13.41 ± 0.25 | 13.33 ± 0.25 |
|  | 26 | 10.45 ± 0.18 | 10.35 ± 0.20 | 9.32 ± 0.19 | 9.19 ± 0.17 |
|  | 32 | 10.11 ± 0.32 | 6.98 ± 0.13 | 6.78 ± 0.13 | 7.19 ± 0.13 |
|  | 34 | 8.26 ± 0.19 | 7.04 ± 0.14 | 6.67 ± 0.15 | 7.87 ± 0.20 |
| Juvenile mortality rate<br>(individual<sup>-1</sup> day<sup>-1</sup>) | 22 | 0.05 ± 0.01 | 0.04 ± 0.01 | 0.02 ± 0.00 | 0.03 ± 0.01 |
|  | 26 | 0.06 ± 0.01 | 0.02 ± 0.01 | 0.04 ± 0.01 | 0.03 ± 0.01 |
|  | 32 | 0.09 ± 0.01 | 0.04 ± 0.01 | 0.05 ± 0.01 | 0.05 ± 0.01 |
|  | 34 | 0.14 ± 0.01 | 0.05 ± 0.01 | 0.09 ± 0.01 | 0.10 ± 0.01 |
|  | 36 | 0.18 ± 0.02 | 0.17 ± 0.02 | 0.18 ± 0.02 | 0.27 ± 0.03 |
| Adult lifespan (days) | 22 | 8.00 ± 0.70 | 7.50 ± 0.54 | 10.04 ± 0.43 | 11.24 ± 0.45 |
|  | 26 | 4.54 ± 0.39 | 9.39 ± 0.45 | 11.51 ± 0.48 | 11.65 ± 0.43 |
|  | 32 | 5.21 ± 0.72 | 8.53 ± 0.43 | 8.66 ± 0.44 | 9.29 ± 0.43 |
|  | 34 | 3.76 ± 0.54 | 5.62 ± 0.44 | 6.09 ± 0.53 | 4.68 ± 0.59 |
| Body size<br>(dry mass (mg)) | 22 | 0.30 ± 0.01 | 0.31 ± 0.01 | 0.44 ± 0.01 | 0.52 ± 0.01 |
|  | 26 | 0.23 ± 0.01 | 0.36 ± 0.01 | 0.50 ± 0.01 | 0.58 ± 0.01 |
|  | 32 | 0.19 ± 0.01 | 0.34 ± 0.01 | 0.38 ± 0.01 | 0.36 ± 0.01 |
|  | 34 | 0.17 ± 0.02 | 0.30 ± 0.01 | 0.36 ± 0.01 | 0.26 ± 0.01 |
| Daily fecundity rate<br>(eggs individual<sup>-1</sup> day<sup>-1</sup>) | 22 | 1.77 ± 0.06 | 1.95 ± 0.05 | 1.99 ± 0.06 | 2.59 ± 0.54 |
|  | 26 | 1.52 ± 0.07 | 1.96 ± 0.08 | 2.54 ± 0.37 | 3.00 ± 0.30 |
|  | 32 | 3.79 ± 0.26 | 3.04 ± 0.33 | 3.55 ± 0.51 | 2.64 ± 0.22 |
|  | 34 | 6.87 ± 1.87 | 5.07 ± 0.82 | 4.91 ± 0.52 | 5.76 ± 0.89 |
| Population-level fitness ($r_m$) | 22 | -0.05 ± 0.03 | 0.06 ± 0.02 | 0.10 ± 0.01 | 0.08 ± 0.02 |
|  | 26 | 0.05 ± 0.02 | 0.12 ± 0.02 | 0.14 ± 0.02 | 0.16 ± 0.02 |
|  | 32 | -0.04 ± 0.04 | 0.24 ± 0.03 | 0.23 ± 0.03 | 0.20 ± 0.03 |
|  | 34 | -0.21 ± 0.05 | 0.22 ± 0.03 | 0.15 ± 0.04 | 0.02 ± 0.05 |
|  | 36 | -2.10 ± 0.15 | -2.11 ± 0.21 | -2.19 ± 0.22 | -2.68 ± 0.20 |

Resource depletion at our lowest resource level (0.183 mg ml^−1^) increased the negative effect of increased temperature on juvenile mortality rate (figure 1a, table 2). As temperatures increased from 22 to 34°C, non-overlapping 95% credible intervals indicate that juvenile mortality rate was significantly higher at low resource concentrations than at intermediate resource concentrations (0.367 mg ml^−1^). At 0.183 mg ml^−1^, it increased from 0.05 at 22°C to 0.14 individual^−1^ day^−1^ at 34°C, whereas at 0.367 mg ml^−1^, it increased from 0.04 to 0.05 individual^−1^ day^−1^ across this temperature range.

Development time varied significantly with the interaction between temperature and resource concentration (ANOVA; *F*_9, 2.24_ = 13.44, *p*<0.001, table 1). Development time decreased with temperature at all resource levels, but the decrease with temperature was greater at the low resource level than at higher resource levels due to resource depletion (figure 1b). At 0.183 mg ml^−1^, development time decreased from 18.30 days at 22°C to 8.26 days at 34°C. Development time at the higher resource levels decreased from approximately 13.50 days at 22°C to ~7.50 days at 34°C (table 2).

Resource depletion at low resource concentrations (0.183 mg ml^−1^) resulted in significant variation in size at maturity (mass, mg) between resource levels (ANOVA; *F*_9,0.92_ = 24.26, *p*<0.001, table 1). Adult size decreased both at warmer temperatures and at low resource concentrations, though the decrease with temperature was greater at higher resource levels than at the low resource level. At low resource concentration, size decreased by 0.13 mg as temperatures increased from 22 to 34°C, while at the highest resource concentration (0.733 mg ml^−1^), size decreased by 0.26 mg (figure 1c, table 2).

Adult lifespan varied significantly with the interaction between temperature and resource concentration (ANOVA; *F*_9, 699.60_ = 7.96, *p*<0.001, table 1). Lifespan was greatest at 0.733 mg ml^−1^, where it was 11.24 days at 22°C, 11.65 days at 26°C, and 4.68 days at 34°C. In contrast, at low resource concentrations, lifespan decreased from 8.00 days at 22°C to 3.76 days at 34°C mg (figure 1d, table 2).

At all resource levels, predicted daily fecundity rate increased with temperature (table 2), though the increase was greater at low resource concentration than at higher resource levels. At low resource concentration, fecundity increased with temperature from 1.77 eggs individual^−1^ day^−1^ at 22°C to 6.87 eggs individual^−1^ day^−1^ at 34°C. At the higher resource levels, fecundity increased from ~2 eggs at 22°C to ~5 eggs individual^−1^ day^−1^ at 34°C.

### Population fitness

At all resource levels, *r*_m_ responded unimodally to temperature. However, resource depletion at low resource concentrations (0.183 mg ml^−1^) significantly depressed *r*_m_ across the entire temperature range (figure 2a) and caused it to peak at a significantly lower temperature than at the intermediate resource level (0.367 mg ml^−1^; figure 2b, table 3). Resource depletion at 0.183 mg ml^−1^ also significantly narrowed the thermal niche width for *r*_m_ compared to higher resource levels (figure 2a, table 3).

**Figure 2.**
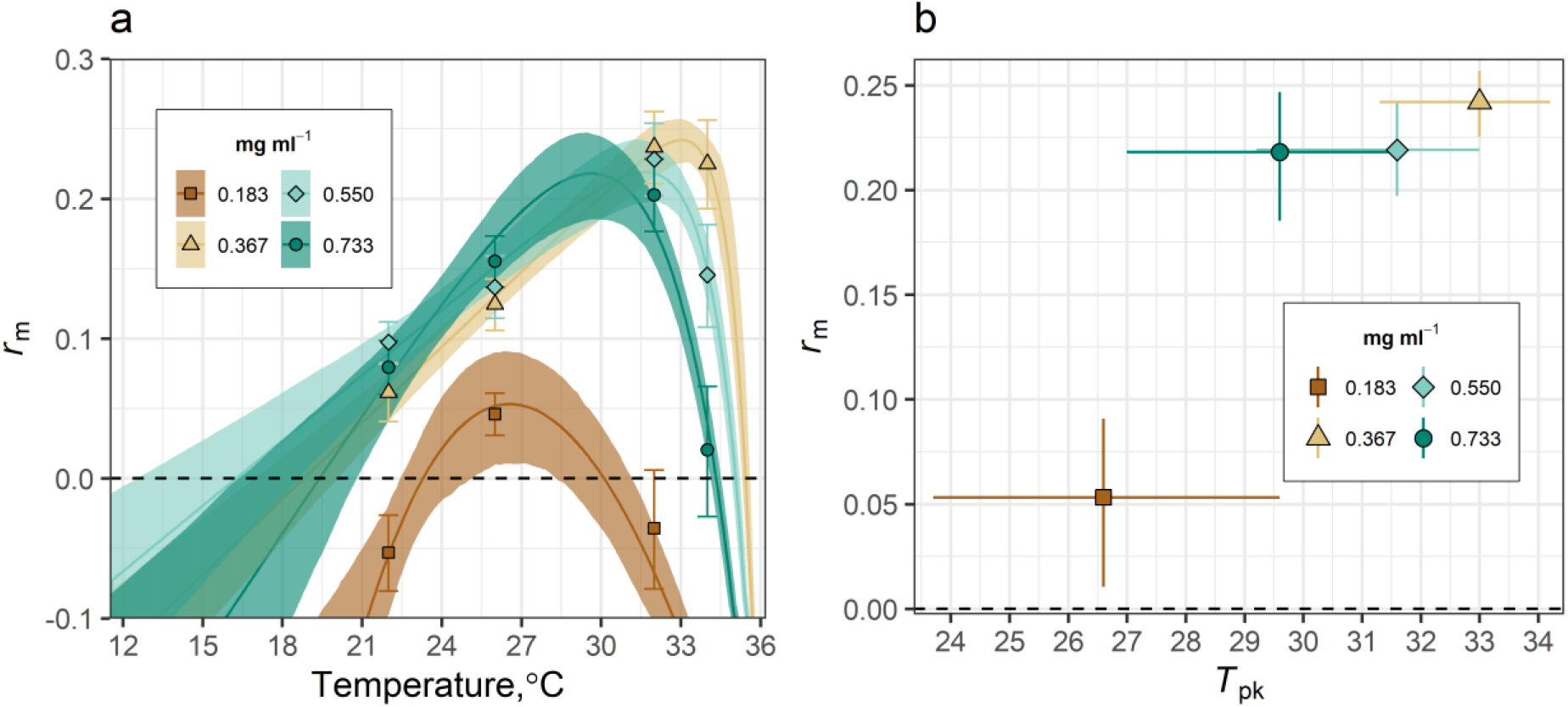
The effect of resource depletion on the thermal response of population-level *Ae. aegypti* fitness (*r*_m_) with bootstrapped 95% prediction bounds. **a,** Resource depletion at 0.183 mg ml^−1^ significantly depressed *r*_m_ across the entire temperature range and narrowed its thermal niche width compared to the higher resource levels (non-overlapping 95% prediction bounds, table 3). Symbols denote matrix projection estimates with 95% confidence intervals (table 2). **b,** Resource depletion at 0.183 mg ml^−1^ significantly (non-overlapping 95% confidence intervals) lowered maximal *r*_m_ and caused it to peak at a significantly lower temperature than at the intermediate resource level (0.367 mg ml^−1^). Predicted *r*_m_ *T*_opt_ at 0.183 mg ml^−1^ indicates that resource depletion could decrease *r*_m_ *T*_opt_ by 6.4°C, when compared to the intermediate resource level (0.367 mg ml^−1^, table 3).

**Table 3.** Parameter estimates of the Thermal Performance Curves of *r*_m_ by resource level. Non-overlapping 95% Confidence Intervals (CIs) indicate that resource depletion at the lowest resource level (0.183 mg ml^−1^) significantly depressed maximal growth (*r*_m_ at *T*_opt_) compared to the higher resource levels. Resource depletion at 0.183 mg ml^−1^ caused a significant decrease in *r*_m_ *T*_opt_ compared to *r*_m_ *T*_opt_ at 0.367 mg ml^−1^. Resource depletion at 0.183 mg ml^−1^ also caused a significantly narrower thermal niche width compared to the higher resource levels.

| Resource concentration<br>(mg ml<sup>-1</sup>) | $r_m$ at $T_{opt}$<br>(± 95% CI) | $T_{opt}$ (°C)<br>(95% CI) | $T_{min}$ (°C)<br>(95% CI) | $T_{max}$ (°C)<br>(95% CI) | Thermal niche width (°C)<br>(95% CI) |
| --- | --- | --- | --- | --- | --- |
| 0.183 | 0.05 ± 0.04 | 26.6<br>(23.7 - 29.6) | 23.3<br>(22.4 - 24.9) | 30.1<br>(28.6 - 31.2) | 6.8<br>(3.7 - 8.8) |
| 0.367 | 0.24 ± 0.02 | 33.0<br>(31.3 - 34.2) | 18.8<br>(17.1 - 20.2) | 35.4<br>(35.4 - 35.7) | 16.6<br>(15.2 - 18.6) |
| 0.550 | 0.22 ± 0.02 | 31.6<br>(29.2 - 33) | 16.2<br>(12.4 - 18.6) | 35.1<br>(35.0 - 35.3) | 18.8<br>(16.4 - 22.9) |
| 0.733 | 0.22 ± 0.03 | 29.6<br>(27.0 - 31.5) | 19.4<br>(16.6 - 21.0) | 34.3<br>(34.2 - 34.6) | 14.9<br>(13.2 - 18) |

At low resource concentration, *r*_m_ was negative until temperatures increased to 23.3°C (figure 2, table 3). At this resource level, *r*_m_ reached a peak of 0.05 at its *T*_opt_ (26.6°C); it then declined to negative growth at 30.1°C. The breadth of *r*_m_’s thermal niche width at the lowest resource concentration was 6.8°C. In contrast, at the intermediate food level (0.367 mg ml^−1^), *r*_m_ became positive as temperatures increased to 18.8°C; it was maximal at 33.0°C (0.24, figure 2, table 3). At 0.367 mg ml^−1^, *r*_m_ declined to negative growth at 35.4°C. The thermal niche width for *r*_m_ at this resource level was 16.6°C. Overlapping CIs indicate that the predicted differences between the intermediate resource level and the higher resource levels (0.550 and 0.733 mg ml^−1^) in *r*_m_ at *T*_opt_, *T*_opt_, and the thermal niche width were non-significant (figure 2, table 3).

### Sensitivity analyses

#### Elasticities

Juvenile traits (development time and survival) were the most important contributors to *r*_m_ (figure 3). For example, at the lowest resource concentration (0.183 mg ml^−1^) at 26°C, a 0.5 proportional increase in juvenile traits would increase rate of increase from 0.046 to 0.063 (figure 3d). In contrast, for the same treatment, a proportional increase of the same magnitude for adult survival would increase *r*_m_ from 0.046 to 0.050 (figure 3e), and fecundity would increase *r*_m_ from 0.046 to 0.048 (figure 3f). This underlines how the temperature-dependence of *r*_m_ derives mainly from how resource depletion impacts juvenile mortality and development, which determine the number of reproducing individuals and the timing of reproduction, respectively. Fecundity and adult survival, on the other hand, have relatively negligible effects on *r*_m_, which suggests that the carry over effect of reduced size at maturity on *r*_m_ is relatively weak.

**Figure 3.**
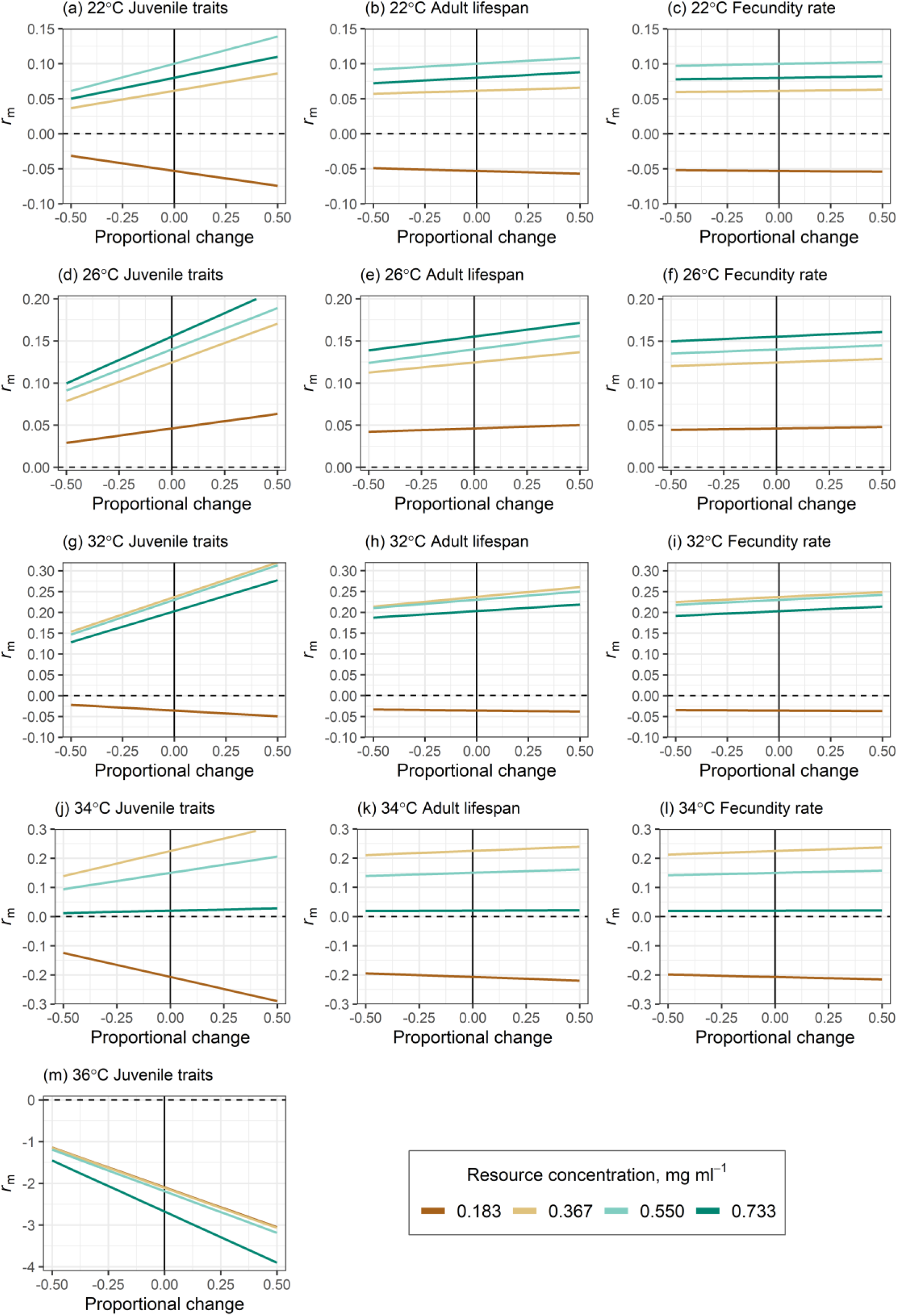
Sensitivity of *r*_m_ to proportional changes in juvenile and adult traits by temperature across resource levels. Juvenile survival and development were the most important contributors to *r*_m_, as relatively small changes in the summed matrix elements for this trait would result in relatively large changes in *r*_m_. Sensitivity of *r*_m_ to adult traits was much weaker compared to sensitivity to juvenile traits.

#### Fecundity estimates

Figure 4 shows that the *r*_m_ TPCs were insensitive to uncertainty in our fecundity estimates. Comparison with the central estimates shows that, for all resource levels, using the upper and lower 95% exponents (Supplementary file 1‒‒Equation S1 and figure S1b) for the scaling between lifetime fecundity and size does not qualitatively change the predicted *r*_m_ TPCs, or the matrix projection *r*_m_ estimates that were used to fit the *r*_m_ TPCs. Predicted *r*_m_ *T*_pk_ was also insensitive to uncertainty in our fecundity estimates. Also, using the upper and lower 95% exponents (Supplementary file 1‒‒Equation S1 and figure S1b) for the scaling between lifetime fecundity and size does not qualitatively change predicted maximal *r*_m_ or *r*_m_ *T*_pk_.

**Figure 4.**
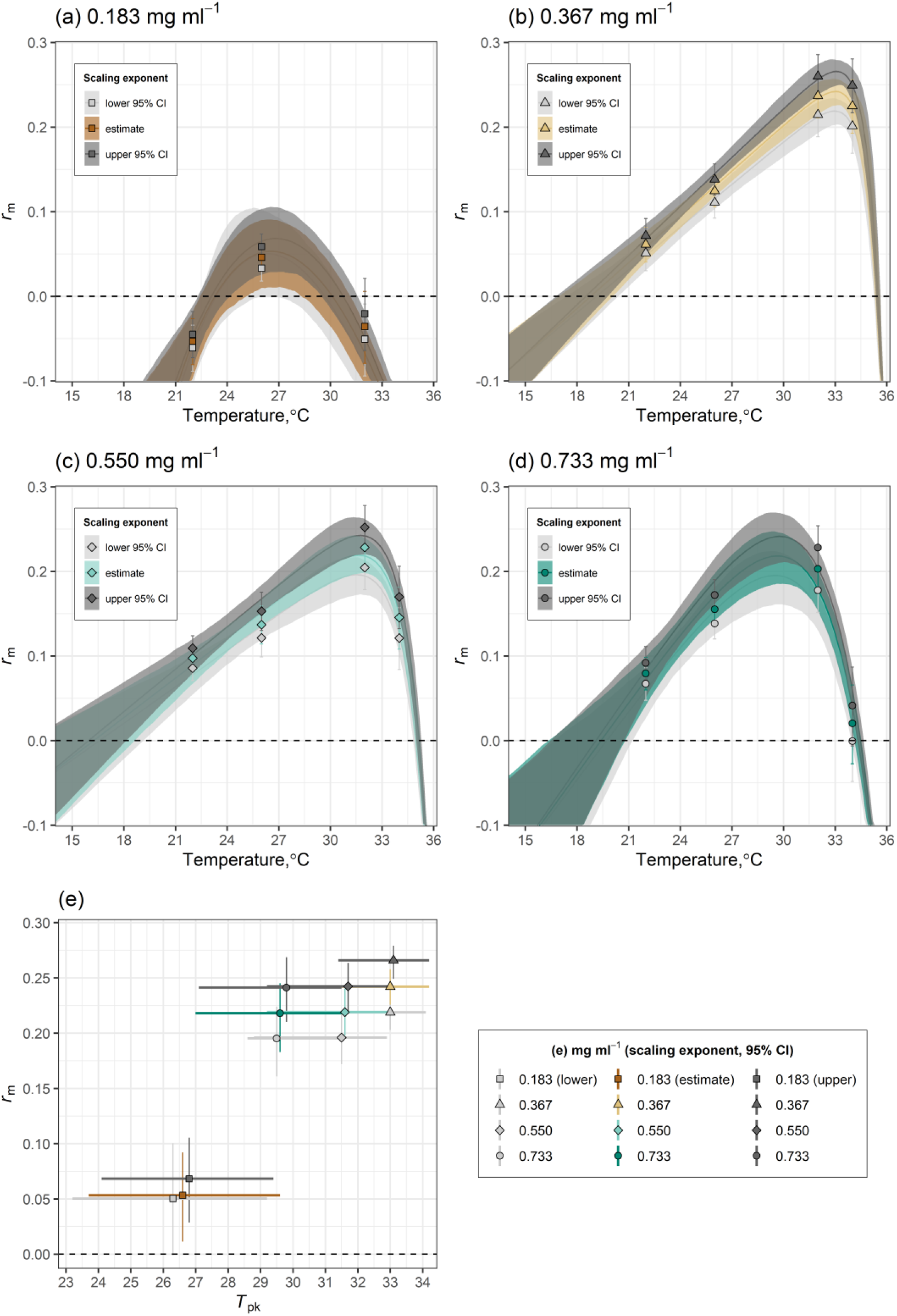
a-d, The insensitivity of the *r*_m_ TPCs to uncertainty in our fecundity estimates by resource level. Comparison with the central estimates (coloured lines and 95% confidence bounds compared with greyscale lines and CBs) shows that, for all resource levels, using the upper and lower 95% exponents (Supplementary file 1‒‒Equation S1, figure S1b) for the scaling between lifetime fecundity and size does not qualitatively change the predicted *r*_m_ TPCs, or the matrix projection *r*_m_ estimates (represented by symbols) that were used to fit the *r*_m_ TPCs. **e, The insensitivity of predicted *r*_m_ *T*_pk_ to uncertainty in our fecundity estimates by resource level.** Using the upper and lower 95% exponents (Supplementary file 1‒‒Equation S1, figure S1b) for the scaling between lifetime fecundity and size does not qualitatively change predicted maximal *r*_m_ (symbols with 95% CIs (vertical coloured lines)) or *r*_m_ *T*_pk_ (symbols with 95% CIs (horizonal coloured lines)).

## Discussion

Our results show that resource depletion in the juvenile habitat can have a profound effect on the temperature dependence of population fitness in *Aedes aegypti*. At the lowest resource level (0.183 mg ml^−1^), resource depletion had a consistent negative effect on the thermal responses of underlying fitness traits (figure 1), which caused a marked divergence between the *r*_m_ TPCs (figure 2). Resource depletion at the lowest resource level significantly depressed *r*_m_ across the entire temperature range, caused a significant decrease in *r*_m_ *T*_opt_ compared to the intermediate resource level (0.367 mg ml^−1^), and significantly narrowed the thermal niche width compared to the higher resource levels (figure 2, table 3). So far, most studies that have used mechanistic trait-based approaches to understand the population-level effects of temperature × resource interactions have focused on aquatic organisms (Orcutt & Porter 1984; Thomas *et al.* 2017; Bestion *et al.* 2018). Our study shows that the combined effects of these ubiquitous environmental factors also need to be considered when predicting the effects of global environmental change on disease vector populations, and other terrestrial arthropods.

The elasticity analysis shows that the primary mechanism underlying the divergent temperature dependence of *r*_m_ across resource levels is increased juvenile development time and mortality at low resource concentration (figure 3). The negative effect of low-resource concentration on these traits delayed the onset of reproduction and population-level reproductive output, respectively. This finding‒‒that juvenile traits contribute more to *r*_m_ than adult traits‒‒is consistent with general studies of fitness in organisms with complex lifecycles (Caswell 1978; Kammenga *et al.* 1996; Huey & Berrigan 2001; Cator *et al.* 2020), including mosquitoes (Juliano 1998; Huxley *et al.* 2020).

Furthermore, because individual fecundity rate and adult lifespan had negligible effects on *r*_m_ compared to juvenile traits, this finding suggests that the carry over effect of reduced size at maturity on *r*_m_ is relatively weak (figure 3). For example, at low-resource concentration, lifetime fecundity was greater at 22°C than at 26°C because body size and adult lifespan were greater at 22°C. Despite this difference, *r*_m_ at 26°C was predicted to be ~200% greater than at 22°C (figures 1 and 2, table 2). This result derives from how juvenile development time almost halved as temperatures increased from 22 to 26°C (table 2). Although juvenile mortality rates for these treatments were similar (0.05 at 22°C versus 0.06 at 26°C, table 2), faster development at 26°C meant that greater numbers of individuals could contribute to population growth through reproductive output. This finding is key because most projections of how warming will affect disease transmission through its effects on vector abundance are based on laboratory populations reared under optimal resource conditions (e.g., Mordecai *et al.* 2013, 2017; Iwamura *et al.* 2020; Shocket *et al.* 2020). Therefore, such projections may be unreliable because they are likely to underestimate the effect of temperature on juvenile traits and overestimate its effect on adult traits.

The trait-level responses of our higher resource concentration treatments correspond with studies that have synthesised laboratory-derived trait responses to temperature to estimate vector fitness and disease transmission. In these studies, the juvenile development rate of most mosquito vectors is expected to increase from ~0.07 day^−1^ at 22°C to ~0.14 day^−1^ at 32°C (Mordecai *et al.* 2019). In the present study, development rate (1/development time; figure 1b, table 2) increased by a similar margin. For example, development rate at 0.550 mg ml^−1^ increased from 0.07 day^−1^ at 22°C to 0.15 day^−1^ at 32°C. In contrast, at low resource concentration, we found juvenile development rate increased from 0.05 day^−1^ at 22°C to 0.12 day^−1^ at 32°C (figure 1b, table 2). Although these differences in juvenile development rate may appear small, we show that they can have dramatic effects on the temperature dependence of *r*_m_ when combined with the negative impact of resource limitation on juvenile survival (figure 1a, table 2).

Juvenile mortality rate increased significantly with temperature and it was consistently higher at low resource concentration (figure 1a) than at higher resource levels. This is probably because somatic maintenance costs increase with metabolic rate (Kooijman 2000), which cannot be met below a threshold resource level. Intensified competition at low resource levels is also likely to have contributed to preventing some individuals from meeting this increased energy demand. This explains why juvenile mortality rates were highest at 32 and 34°C at low-resource concentration (except at 36°C where no individuals survived at all) where the energy supply-demand deficit was expected to be the largest.

Since resource depletion can mediate the temperature dependence of *r*_m_, it is also important to determine the temperature dependence of resource availability itself (Huey & Kingsolver 2019). For example, the natural diet of mosquito larvae comprises of detritus and microbial decomposers (Merritt *et al.* 1992), which are both sensitive to temperature (Craine *et al.* 2010; Smith *et al.* 2019). Therefore, shifts in environmental temperature could alter the concentration of food in the environment, which could affect the growth of detritivore populations. While recent studies have provided useful insights into the relationships between microbes, detritus and mosquito vectors (Yee *et al.* 2007; Chouaia *et al.* 2012; Dickson *et al.* 2017; Souza *et al.* 2019; Hery *et al.* 2021), future work could focus on the temperature-dependencies of these relationships.

Such a focus could provide important insights into how disease vectors and other arthropods will respond to environmental change. For example, if resource availability increases with climatic warming (e.g., due to increases in decomposition and microbial growth rates), its regulatory effect on population growth and abundance could be relaxed through increased juvenile development and adult recruitment rates. Indeed, increased resource availability with warming could contribute to the expansion of disease vectors and other invasive pest species into regions that were previously prohibitive by broadening *r*_m_’s thermal niche width (Amarasekare & Simon 2020; Lehmann *et al.* 2020). On the other hand, evidence from our high resource concentration treatments (e.g., a lower *T*_opt_ at 0.733 than at 0.367 mg ml^−1^) may suggest that warming could have a negative impact on population growth by causing resources to be overabundant, which could lead to eutrophication and hypoxia in aquatic environments (Liikanen *et al.* 2002).

Alternatively, if climate change reduces resource availability (e.g., by disrupting temperature-dependent consumer-resource relationships), species’ spatiotemporal ranges could contract (Huey and Kingsolver 2019, Lister and Garcia 2018). This is because, as we have shown here, resource depletion at low food levels can prevent *r*_m_ from being positive at lower temperatures, can lower *r*_m_ *T*_opt_, and can force *r*_m_ to become negative at lower temperatures. In this way, the effects of rising temperatures on vulnerable arthropod populations could be especially pernicious, if resource availability is simultaneously reduced (Huey & Kingsolver 2019).

We did not measure the effect of temperature and resource concentration on fecundity directly, but used the size-scaling of this trait to estimate this effect. This is because most of the effect of intensified larval competition at low resource concentration, is expected to affect adult mosquitoes indirectly by reducing size at emergence and lifespan (Briegel 1990, Steinwascher 1982). Despite these assumptions, we show that substantial under- or overestimation of fecundity by our size-scaling predictions and the use of starved adult lifespans, would not affect our main conclusions. This is because predicted fitness was relatively insensitive to these traits (figures 3 and 4).

Although the increased negative carry over effects of temperature at low resource levels on adult traits may have had a relatively weak impact on fitness compared to juvenile traits, temperature × resource interactions may have important effects on other components of vector-borne disease transmission (Parham *et al.* 2015). For example, body size is expected to affect vectorial capacity because larger individuals are more likely to outlive a pathogen’s extrinsic incubation period (Nasci 1986; Ameneshewa & Service 1996). However, evidence shows that temperature and resource availability can act interactively to influence the relationships between body size, longevity and vector competence (Barreaux *et al.* 2016, 2018). Indeed, as we have shown here, resource limitation can exaggerate the negative relationship between size and temperature (Atkinson 1994). This effect could increase transmission probability as smaller *Ae. aegypti* may compensate for poor larval nutrition by biting more frequently (Scott *et al.* 2000). Also, while studies of the independent effects resource availability and temperature in the juvenile stages on within-vector pathogen development report mixed results (Kay *et al.* 1989; Dodson *et al.* 2012; Shapiro *et al.* 2016, 2017), future studies could consider how the combined effects of temperature and resource availability affect this, and other important transmission traits.

Another important elaboration of this study’s findings for vector-borne disease research is that it highlights the need to develop realistic and tractable methods of measuring density-dependent effects on population fitness in the field. Without such datasets it will not be possible to link temperature- and resource-dependent fitness to vector abundance dynamics and VBD dynamics. Semi-field systems offer a way to track the entire vector lifecycle under ambient environmental conditions (Jones *et al.* 2021). Such systems could allow for the effects of temperature × resource interactions on fitness and abundance to be explored under conditions which more closely resemble natural environments. For example, the effects larval competition could be studied more directly by varying the number of larvae across gradients of temperature and resource availability. Indeed, variation in larval density may introduce additional fitness constraints through interference, exploitative competition and the accumulation of waste products.

Rapid global change is expected to have far-reaching and disruptive ecological impacts (Trisos *et al.* 2020). Climate-driven shifts in the spatiotemporal distributions and abundances of organisms are likely to cause widespread harm to ecosystems, biodiversity and society (Parmesan 2006; Diagne *et al.* 2021). This concern has prompted calls for a more complete understanding of how interactions between environmental factors can affect population-level responses (Cross *et al.* 2015; Huey & Kingsolver 2019; Taheri *et al.* 2021). So far, most studies that have used mechanistic approaches to understand the population-level effects of temperature × resource interactions have focused on aquatic organisms (Orcutt & Porter 1984; Thomas *et al.* 2017; Bestion *et al.* 2018). However, our study provides compelling evidence that the combined effects of these ubiquitous environmental factors also need to be considered when predicting the effects of global environmental change on disease vector populations, and other terrestrial arthropods.

## Materials and Methods

To investigate the effects of temperature and resource depletion on mosquito life history, we employed a 5×4 factorial design comprised of five temperatures (22, 26, 32, 34, and 36°C) and four resource concentration levels (0.183, 0.367, 0.550 and 0.733 mg ml^−1^). These experimental temperatures span the range of temperatures that this strain of *Ae. aegypti* (F16-19 originating from Fort Meyer, FL; Bargielowski *et al.* 2013) is likely to experience in the wild between May (the onset of mosquito season) and November (Arguez *et al.* 2012). We extended our range to 36°C to determine the upper critical thermal limit for this strain. Our resource concentration levels are within the range of studies that have investigated the effects of depleting juvenile resource environments on *Aedes aegypti* (Subra & Mouchet 1984). Our lowest resource level (0.183 mg ml^−1^) was chosen to simulate a level of resource limitation that is expected in natural juvenile habitats (Arrivillaga & Barrera 2004; Barrera *et al.* 2006). Further, our preliminary assays showed that resource levels below 0.183 mg ml^−1^ resulted in complete juvenile mortality.

The experiment was carried out in two blocks with treatments randomly assigned to one of these blocks. On Day 0, batches of approximately 800 eggs were deposited into five (one per experimental temperature) plastic tubs containing 300 ml of dechlorinated tap water. Each tub was provided with a pinch of powdered fish food (Cichlid Gold®, Hikari, Kyrin Food Industries Ltd., Japan) to stimulate overnight hatching. Tubs were randomly assigned to a water bath (Grant Instruments: JAB Academy) set at one of the five experimental temperatures. Water baths were situated in a 20°C climate-controlled insectary with a 12L:12D photoperiod and 30 minutes of gradual transition of light levels to simulate sunrise and sunset. On the following day (Day 1), we created the treatments by separating first instar larvae were into cohorts of 50, which were then transferred to clean tubs containing 300 ml of fresh water. Each treatment comprised of three replicate tubs (3×50 individuals treatment^−1^). Resource concentration levels were attained by adding 55, 110, 165 and 220 mg of powdered fish food to the tubs, respectively. To allow resource depletion, tubs received two pulses of equal quantity. Half of the assigned quantity was provided on Day 1; the remaining half was provided on Day 4. After Day 4, resource levels were not adjusted but water volumes were topped up, if necessary.

### Fitness calculation

To calculate *r*_m_, we used our life history trait data to parameterise stage-structured matrix projection models (Equation 1; Caswell 1989), which describe change in a population over time:

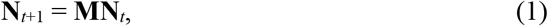

where **N**_*t*_ is a vector of abundances in the stage classes at time *t* and **M** is the population projection matrix. The first row of **M** is populated by daily fecundity (the number of female offspring produced per female at age *i*). The sub-diagonal of **M** is populated with the survival proportions from age *i* to age *i*+1. Multiplying the transition matrix (**M**; Equation 1) and stage-structured population size vector (**N**_*t*_; Equation 1) sequentially across time intervals yields the stage-structured population dynamics. Once the stable stage distribution of the abundance vector is reached, the dominant eigenvalue of the system is the finite population rate of increase (*λ*) (Caswell 1989). Then, the intrinsic rate of population growth is

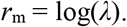

This is a population’s inherent capacity to reproduce, and therefore a measure of population-level fitness (Birch 1948; Cole 1954; Savage *et al.* 2004). Negative *r*_m_ values indicate decline and positive ones, growth. The projection matrices were built and analysed using the `popbio` R package (Stubben & Milligan 2007; R Core Team 2018).

### Model parameterisation

#### Immature development time and immature and adult survival proportions

Matrix survival elements (the sub-diagonal of the matrix **M**; Equation 1) were populated with continuous survival proportions estimated using the Kaplan-Meier survival function in the survival R package (Therneau 2021). We assumed life stage duration (i.e. larva-to-pupa-to-adult) was the mean duration of transitioning into and out of that stage, and a fixed age of adult emergence at the mean age of emergence. Adult survival elements were populated with the Kaplan-Meier proportions. Hatching-to-adult development times were calculated by recording the day and time that egg eclosion, pupation and adult emergence occurred for each individual. Upon pupation, mosquitoes were held in individual falcon tubes containing 5 ml of tap water. This enabled pupa-to-adult development durations and the lifespans of individual starved adults to be recorded. Adult lifespan was recorded in the absence of food, which forces adults to metabolise nutritional reserves accumulated during larval development. Therefore, starved adult lifespan should increase with size at emergence, so it is a useful indicator of the carry over effects of temperature and resource availability in the larval habitat (Briegel 1990; Agnew *et al.* 2002).

#### Daily fecundity rate

Fecundity and body size are positively related in many insect taxa, including mosquitoes (Honěk 1993). For this reason, scaling relationships between fecundity and size are commonly used in predictions of population growth in *Aedes* (Livdahl & Sugihara 1984; reviewed in Juliano & Lounibos 2005). A detailed description of our method for estimating fecundity is provided in Supplementary file 1. Briefly, we measured individual dry mass, and estimated lifetime fecundity using previously published datasets on the temperature-dependent scaling between mass and wing length (van den Heuvel 1963), and wing length and fecundity (Briegel 1990; Farjana & Tuno 2012). Daily fecundity rate is required for the matrix projection models (Equation 1), so we divided lifetime fecundity by lifespan and multiplied by 0.5 (assuming a 1:1 male-to-female offspring ratio) to give temperature-specific individual daily fecundity. Later, we show that this much variation in the scaling of fecundity does not qualitatively change our results.

### Parameter sensitivity

We used the delta method to approximate 95% confidence intervals (CIs) to account for how uncertainty in survival and fecundity estimates is propagated through to the *r*_m_ estimate (Caswell 1989; Skalski *et al.* 2007). This method requires the standard errors of the survival and fecundity element estimates. For survival, we used the standard errors estimated by the Kaplan-Meier survival function in the `survival` R package. For fecundity, we calculated the standard errors of the mean daily fecundity rates (Supplementary file 1‒‒table S2) for each treatment using the `Rmisc` R package (Hope 2013). As an additional sensitivity analysis, we recalculated fitness using the upper and lower 95% CIs of the exponents for the scaling of size and lifetime fecundity (figure 3).

### Elasticity analysis

We used elasticities to quantify the relative contributions of individual life history traits to *r*_m_. Elasticity, *e_ij_*, measures the proportional effect on *λ* of an infinitesimal change in an element of **M**(Equation 1) with all other elements held constant (the partial derivative) (Caswell *et al.* 1984; de Kroon *et al.* 1986). This partial derivative of *λ*, with respect to each element of **M**, is *s_ij_* = ∂*λ*/∂*a_ij_* = *v_i_w_j_* with the dot product 〈**w**, **v**〉 = 1. Here, **w** is the dominant right eigenvector (the stage distribution vector of **M**), **v** is the dominant left eigenvector (the reproductive value vector of **M**), and *a_ij_* is the *i*×*j*^th^ element of **M**. Elasticities can then be calculated using the relationship: *e_ij_* = *a_ij_*/*λ × s_ij_*. Multiplying an elasticity by *λ* gives the absolute contribution of its corresponding *a_ij_* to *λ* (Caswell *et al.* 1984; de Kroon *et al.* 1986). Absolute contributions for juvenile and adult elements were summed and changed proportionally to quantify the sensitivity of *r*_m_ to these traits.

### Statistical analyses

All statistical analyses were conducted using R (R Core Team 2018).The trait data (adult lifespan and body size) were normally distributed, so we used full factorial linear models (LM) to determine the significance of each predictor on the thermal response of each of these traits. The development time data were not normally distributed, so we used a generalized linear model (GLM) with gamma distribution and identity link functions (predictor effects were considered additive). Replicate was included in all regression models as a fixed effect. We investigated the effect of resource level on the temperature dependence of daily per capita juvenile mortality rate by fitting an exponential function to the survival data with R package `flexsurv` (Jackson 2016). The final mortality model was obtained by dropping terms from the full model (consisting of temperature × resource level + replicate as fixed effect predictors). If removing a term worsened model fit (ΔAIC > ‒2), then it was retained. Otherwise, it was removed (Supplementary file 1‒‒table 1). Maximum likelihood methods executed in `flexsurv` were used to estimate the juvenile mortality rate (and their 95% CIs) for each treatment. These estimates were then used to determine the significance of the effects of temperature and resource depletion on juvenile mortality.

### Quantifying *r*_m*’s*_ thermal performance curve

To determine how resource depletion affected the shape of the *r*_m_ TPC, we fitted several mathematical models that allow for negative values at both cold and hot extremes, including polynomial models using linear regression, as well as non-linear models with non-linear least squares (NLLS) using the `rTPC` R package (Padfield *et al.* 2020). Overall, the Lactin2 (Lactin *et al.* 1995) and Kamykowski (Kamykowski & McCollum 1986) models were equally best-fitting according to the Akaike Information Criterion (AIC) (Supplementary file 1‒‒table 2). From these, we picked the Kamykowski model because it was better at describing the estimated *r*_m_ at our lowest resource level. This model is defined as

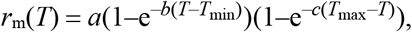

where *T* (°C), and *T*_max_ and *T*_min_ are the high and low temperatures at which *r*_m_ becomes negative, respectively, and *a*, *b*, and *c*, are shape parameters without any biological meaning. Bootstrapping was used to calculate 95% prediction bounds for each *r*_m_ TPC (Padfield *et al.* 2021) and confidence intervals (CIs) around its *T*_opt_, as well as the thermal niche width (*T*_max_ ‒ *T*_min_).

## Acknowledgements

All authors contributed to the conception of the study and designed the experiments, L.J.C. provided the mosquitoes; P.J.H. and S.P. performed the modelling; P.J.H. collected the data and analysed it. P.J.H. wrote the first draft of the manuscript, and all authors contributed substantially to revisions.

