## Supplementary file 1 for "The effect of resource depletion on the thermal response of mosquito population fitness"

### The effect of resource depletion on the thermal response of mosquito population fitness

*Method used to predict fecundity rate for matrix projection models (main text Equation 1)*

We measured each individual's dry mass to the nearest 0.01 mg using a microbalance. Prior to weighing, mosquitoes were dried individually in microcentrifuge tubes containing desiccant-silica gel for a minimum of 14 days. For the temperature-dependent scaling between mass and wing length, we analysed the van Heuvel (1963) dataset. This showed that as temperatures increase from 25 to 34°C, the scaling between mass and wing length changes significantly (figure S1a). Our analysis of the Farjana (Farjana & Tuno 2012) dataset indicated that the scaling between wing length and fecundity changes significantly with temperature but not resource level (figure S1b).

To estimate lifetime fecundity ($F$ in Equation S1) from mass for mosquitoes that we reared at 22°C at all food densities, we predicted wing length from mass using the mass-to-wing length exponent at 25°C in the van Heuvel (1963) dataset. We used these wing lengths to predict fecundity using the wing length-to-fecundity scaling exponent from the Farjana (Farjana & Tuno 2012; $n$ = 264, $R^2$ = 0.87, $p$<0.001; Equation S1) dataset at their at 20°C.

For mosquitoes that we reared at 26°C, there is no corresponding temperature treatment in the Farjana dataset (Farjana & Tuno 2012), so we first predicted wing length from mass using the mass-to-wing length exponent at 25°C in the van Heuvel (1963) dataset. We then predicted fecundity using the wing length-to-fecundity scaling from the Briegel (1990) dataset at 27°C ($n$ = 206, $R^2$ = 0.77, $p$<0.001; Equation S1). For mosquitoes that we reared at 32 and 34°C, we predicted wing length from mass using the mass-to-wing length exponent at 34°C in the van Heuvel (1963) dataset. We then predicted fecundity for these mosquitoes using the wing length-to-fecundity scaling exponent from the Farjana (Farjana & Tuno 2012; Equation S1) dataset at 30°C. Fecundity was not estimated at 36°C, as no adults emerged at this temperature. The **scaling equations used to estimate temperature-dependent fecundity from wing length for our mosquitoes are:**

$$\begin{aligned} &22°\text{C}, F = 0.93 + 3.16 \log(L) \\ &26°\text{C}, F = 0.40 + 3.80 \log(L) \\ &32°\text{C}, F = 0.26 + 4.08 \log(L) \\ &34°\text{C}, F = 0.26 + 4.08 \log(L) \end{aligned} \qquad \text{(S1)}$$

The coefficients were derived from our analysis (figure S1) of the Farjana (Farjana & Tuno 2012) and Briegel (1990) datasets.

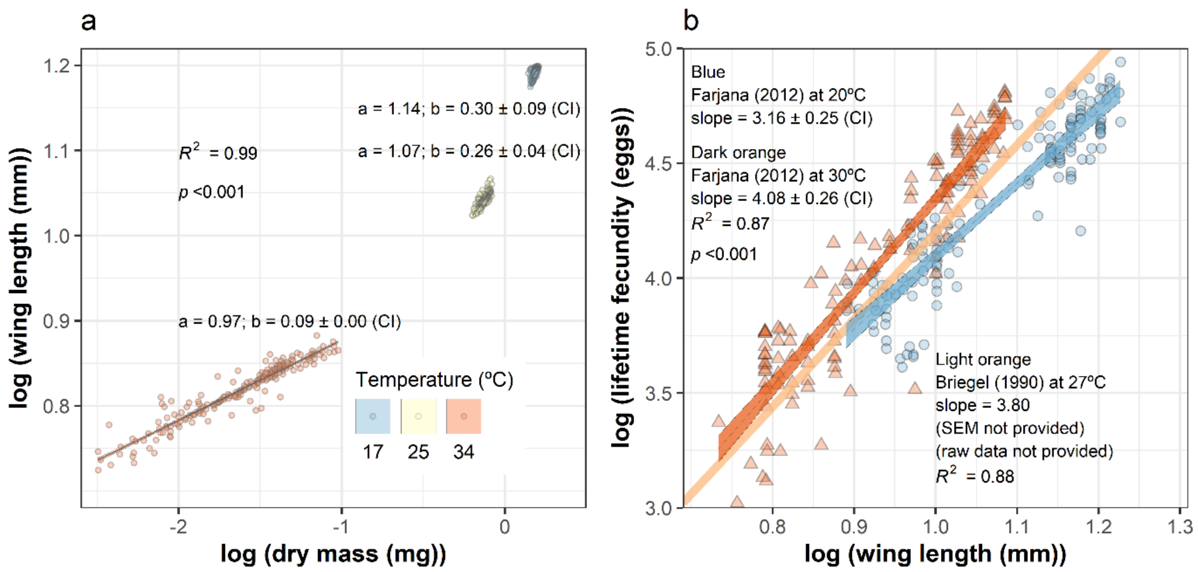


**Figure S1**. **a, Analysis of the van Heuvel** (1963) **dataset shows that the scaling of mass and wing length in *Ae. Aegypti* is temperature-dependent.** The scaling exponents (slopes) for 17°C and 25°C are significantly higher than at 34°C. However, the higher scaling exponent for 17°C is non-significantly higher than for 25°C. **b, Analysis of the Farjana** (Farjana & Tuno 2012) **dataset shows that the scaling of wing length and fecundity in *Ae. Aegypti* is temperature-dependent.** The scaling exponents (slopes) for both resource levels are significantly higher at 30°C than at 20°C. However, the effect of resource on fecundity is non-significant at the temperature level (not shown). The standard error for the scaling exponent at 27°C is not shown because it is not provided in (Briegel 1990), so for 26°C, we assumed a similar 95% CI to those in the Farjana (Farjana & Tuno 2012) dataset (3.80 ± 0.25). Despite these assumptions relating to fecundity, our $r_m$ calculations are robust to uncertainty/variation in the underlying scaling and temperature dependencies (figure 4).

| Model terms | Model name | AIC | ΔAIC | df |
|---|---|---|---|---|
| **Temperature × RC** | **Interaction** | **6446.77** | **0** | **20** |
| Temperature × RC + replicate | Full | 6448.89 | +2.13 | 22 |
| Temperature + RC | No interaction | 6462.33 | +15.56 | 8 |
| Temperature | Temperature only | 6481.32 | +34.55 | 5 |
| Resource | Resource only | 6899.22 | +452.46 | 4 |
| None | Null | 6906.62 | +459.86 | 1 |

**Table S1. Simplification of the exponential juvenile survival model**. The full model includes the effects of temperature × resource concentration (RC) and replicate on mortality. The final mortality model was obtained by dropping terms from the full model (consisting of temperature × resource level + replicate as fixed effect predictors). If removing a term worsened model fit (ΔAIC > –2), then it was retained. Otherwise, it was removed. ΔAICs were calculated as differences from the interaction model (bold).

| Resource concentration (mg ml$^{-1}$) | Model name | AIC | df |
|---|---|---|---|
| 0.183 | Kamykowski (1986) | –44.43 | 10 |
| 0.183 | Lactin2 (1995) | –42.77 | 11 |
| 0.367 | Kamykowski (1986) | –65.53 | 10 |
| 0.367 | Lactin2 (1995) | –67.77 | 11 |
| 0.550 | Kamykowski (1986) | –61.31 | 10 |
| 0.550 | Lactin2 (1995) | –63.61 | 11 |
| 0.733 | Kamykowski (1986) | –53.82 | 10 |
| 0.733 | Lactin2 (1995) | –56.40 | 11 |

**Table S2. Comparison of model fitting for $r_m$ TPCs by resource concentration.** We considered several models that allow for negative values at both cold and hot extremes, including polynomial regression models (quadratic models underfitted the matrix projection $r_m$ estimates, whereas cubic models overfitted these estimates (not shown) and other TPC models (not shown) that are implemented in the `rTPC` R package (Padfield *et al.* 2020). Overall, the Lactin2 function (1995) and Kamykowski model (1986) best described the matrix projection estimates according to the Akaike Information Criterion (AIC). Although these models performed similarly according to their AICs, we chose the Kamykowski model (1986) because it was better at describing the estimated $r_m$ at our lowest resource level.

========================================
